# PLANAtools - an interactive gene expression repository for the planarian *Schmidtea mediterranea*

**DOI:** 10.1101/2022.09.21.508730

**Authors:** Michael Hoffman, Omri Wurtzel

## Abstract

**Motivation:** Planarians are a widespread model for studying regeneration. Major efforts for studying gene function in planarian regeneration produced massive datasets, including transcriptome-wide gene expression analyses from hundreds of conditions. However, the accessibility of gene expression datasets to investigators is limited because of the need for expertise in gene expression analysis in this model, the requirement for computational resources, and the lack of a curated planarian gene expression metadata resource associating samples and their controls.

**Results:** We implemented a computational resource, PLANAtools, that is available online and provides a portal to the analysis of over 160 gene expression analyses. Planarian gene expression datasets from the last decade were processed using a standardized pipeline based on curated planarian metadata. PLANAtools generates plots, annotations, and analyses of gene expression data, based on user parameters.

**Availability:** PLANAtools is implemented using the R/Shiny framework and is accessible from https://wurtzellab.org/planatools

## Introduction

Planarians are highly regenerative flatworms. They can regrow any missing organ by activating injury and regeneration-specific gene expression programs (Reddien, 2018). The planarian genome encodes over 20,000 genes (Rozanski *et al.*, 2019), and therefore understanding the genetic basis for regeneration in this seemingly simple organism is a complex task. The use of RNA-sequencing (RNA-Seq) for gene expression analysis has transformed the study of planarian gene function, by facilitating the collection of thousands of data points in a single experiment. The accessibility of RNA-Seq technologies is demonstrated by the rapid growth in the number of RNA-Seq libraries from planarians, with over 3,450 planarian RNA BioSamples deposited to the publicly available Sequence Read Archive (Leinonen *et al.*, 2011). The availability of transcriptional profiling has driven the discovery of key factors in planarian regeneration and stem cell (neoblast) biology. This includes the discovery of transcription factors associated with lineage selection and maturation (Forsthoefel *et al.*, 2012; Tu *et al.*, 2015), injury response genes (Wenemoser *et al.*, 2012; Wurtzel *et al.*, 2015), and regulators of neoblast differentiation (Zhu *et al.*, 2015; Lapan and Reddien, 2012).

RNA-Seq data allows, in principle, highly reproducible research using common bioinformatic processing tools that are available to the scientific community. Compared to other research modalities, the format of high-throughput sequencing data is standardized, machine readable, and found in accessible repositories (Cock *et al.*, 2010; Leinonen *et al.*, 2011). However, despite these properties, using or comparing RNA-Seq data across projects and research groups presents several challenges. Data processing and analysis differ between scientists and teams, and continuously evolve based on the available tools, protocols, and technology. Moreover, RNA-Seq data is commonly found in raw fastq format, requiring computational proficiency and resources for even basic analysis. Even processed RNA-Seq data, often available as supplementary information in manuscripts can be difficult to use. For example, different transcriptome assemblies are used in the planarian community for RNA-Seq data mapping, limiting comparisons of available data. Major planarian computational resources, such as PlanMine and Planosphere, have become instrumental for the research community by providing powerful tools for studying gene function and genome browsing using a gene or transcript focused interface (Rozanski *et al.*, 2019; Nowotarski *et al.*, 2021). Yet, a major strength of RNA-Seq data is the ability to examine changes in expression of multiple transcripts simultaneously.

Here, we curated published planarian RNA-Seq data and implemented a computational pipeline, based on broadly used RNA-Seq analysis tools for analyzing gene expression. We developed a web-application, PLANAtools, that facilitates browsing, mining, downloading, and visualizing planarian RNA-Seq data interactively, requiring minimal computational skills and resources from the user.

## Results

### Data retrieval and processing

A list of deposited *Schmidtea mediterranea* bulk RNA-Seq metadata was retrieved from the NCBI Sequence Read Archive (Leinonen *et al.*, 2011). The list was curated manually and each library was associated with its controls and replicates. Each set of replicates and their controls was processed as a single biological experiment. Raw RNA-Seq libraries were retrieved from the SRA using the fasterq-dump tool (Leinonen *et al.*, 2011) or by using bowtie2 --sra-acc parameter (Langmead and Salzberg, 2012). Raw RNA-Seq data was mapped to the planarian dd_v6 transcriptome assembly (Rozanski *et al.*, 2019) using bowtie2 with parameter --fast. Paired-end RNA-Seq data was mapped as single-end by considering the first read in a read pair. The mapped data was transformed to Binary Alignment Map (BAM) format using SAMtools (Li *et al.*, 2009). For each biological experiment (i.e., replicates and controls), a read count table was produced using featureCounts v2 with parameters [-M -s 0] (Liao *et al.*, 2014). Data normalization and differential gene expression calling were then performed using DESeq2 using the rlog and DESeqDataSetFromMatrix functions (Love *et al.*, 2014). Differential gene expression analysis was not performed on conditions where no biological replicates were available. Processed data tables were then indexed using the R data.table package and stored as binary files using the saveRDS function.

### Application implementation

PLANAtools is accessible from https://wurtzellab.org/planatools. It was developed in R using the Shiny framework and packaged to a docker image. The current release includes gene expression analyses of 168 assays from over 40 manuscripts. Data browsing was implemented to accomplish four main uses (Fig 1): (1) Showing gene expression in published datasets as a heatmap or volcano plot based on transcripts that are selected by the users. Transcript selection is performed using free text search of transcript IDs or their putative annotation, as previously described (Dagan *et al.*, 2022). (2) Showing differentially expressed genes of an assay as a heatmap, a volcano plot, or a table in the datasets that are available and analyzed with PLANAtools. (3) Presenting gene expression across datasets by comparing the expression of a single gene in control and treatment samples across assays. (4) Finally, to intersect a list of queried genes with a list of differentially expressed genes in an assay. Resulting visualizations are interactive online, and downloadable as vector images for further processing. The processed gene expression tables and differential gene expression analyses are also available for download.

**Figure 1.**
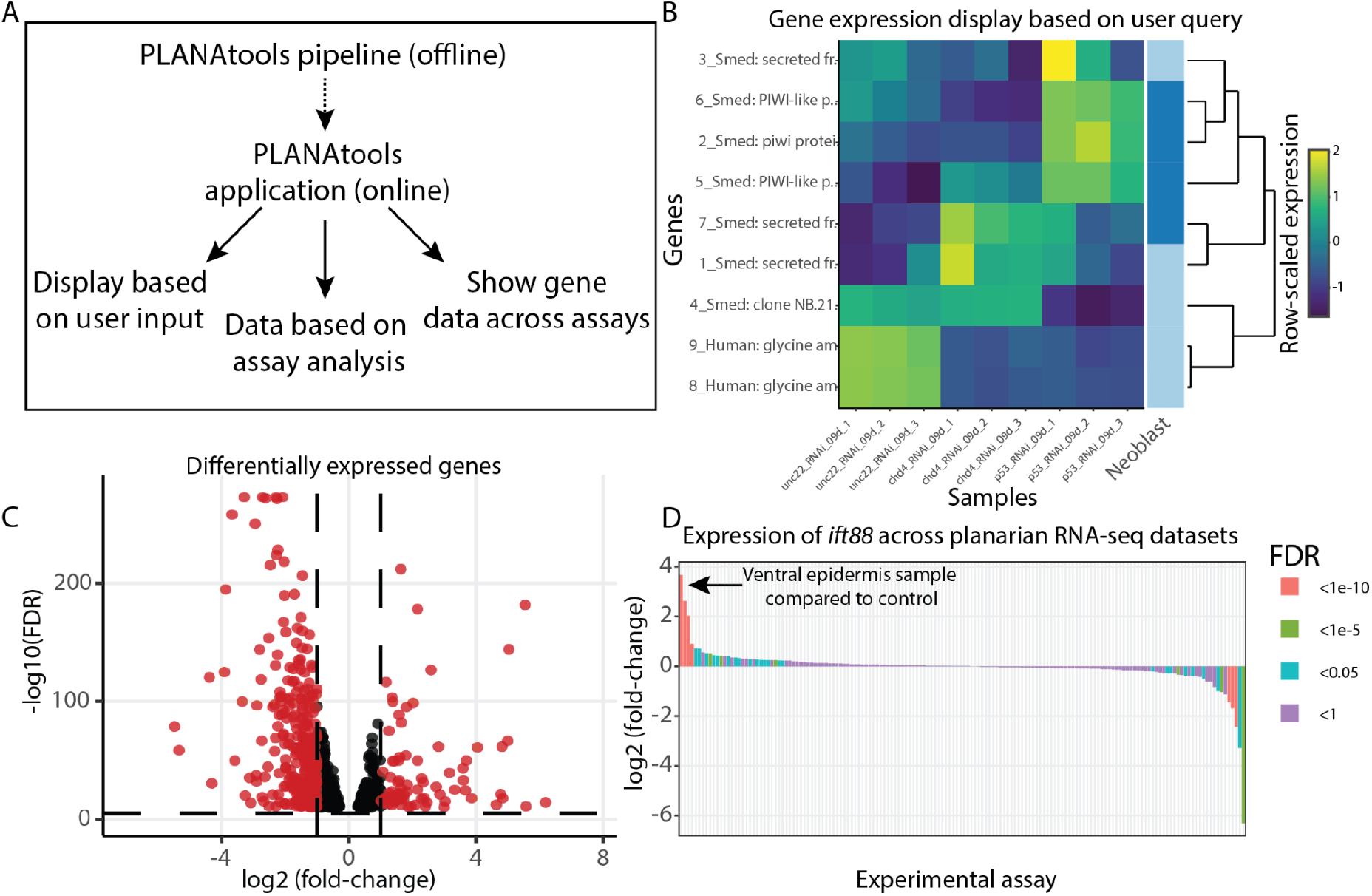
Overview of PLANAtools. (A) Main functions of PLANAtools. An asynchronous standardized pipeline was designed for offline RNA-Seq data processing. The products of this pipeline are used in the online application. Main functions of the online application are shown. (B-D) Images produced by PLANAtools were minimally edited for assembling the multi-panel figure; in the application the plots are interactive and show additional information upon mouse hover. (B) Data visualization produced using the heatmaply package (Galili *et al.*, 2018) on user queried transcripts, using published planarian gene expression data (Tu *et al.*, 2015) that is available on PLANAtools. The right column of the heatmap highlights genes that are known to be expressed in a user selected cell type (Fincher *et al.*, 2018); dark and light blue, expressed or not expressed specifically in the cell type, respectively; in the example shown, the selected cell type is neoblast. (C) A volcano plot showing differentially expressed genes in re-analysis of previously published data (Tu *et al.*, 2015) at 9 days following inhibition of *CHD4* expression. Black and red, significant and insignificant changes to gene expression, based on user parameters, respectively. (D) Differential expression of a user queried gene (*Smed-ift88*) is shown across all available RNA-Seq assays in PLANAtools. Each bar represents a single assay comparing expression in the condition compared to its control. Bar color represents the significance of the differential expression in that assay. The arrow indicates the largest increase in gene expression observed across all assays. The expression of *ift88* is highly enriched in the ventral epidermis compared to its control (Wurtzel *et al.*, 2017).

### Usage

PLANAtools is a hub for gene expression analyses, and facilitates data visualization, browsing and acquisition. In addition, PLANAtools provides links to the original publication, to the raw deposited data in SRA (Leinonen *et al.*, 2011), and to major organism-specific gene resources, such as flybase, planmine, and wormbase (Thurmond *et al.*, 2019; Rozanski *et al.*, 2019; Harris *et al.*, 2020). The functionality of the website is showcased in a video tutorial that is found on the website demonstrating the major features of the application.

### Summary

PLANAtools provides non-experts access to planarian gene expression data, gene annotations, and external data sources. In contrast to most planarian resources, it is not gene- or transcript-centric. Instead, it allows an overview of gene expression using interactive visualizations and direct downloads of the original data. Therefore, it bridges the gaps in the ability to access, use, and re-use planarian gene expression resources.

